# The effect of *in vitro* starch digestibility on glycemic/insulinemic index of biscuits and bread made from non-conventional wholemeal/wholegrain flour mixtures

**DOI:** 10.1101/2023.01.08.523146

**Authors:** Charalampos Papadopoulos, Constantine Anagnostopoulos, Athanasios Zisimopoulos, Maria Panopoulou, Dimitrios Papazoglou, Anastasia Grapsa, Thaleia Tente, Ioannis Tentes

## Abstract

**BACKGROUND:** Carbohydrates as starch are a staple part of the Mediterranean diet. Starch is digested in the small intestine and the resulting glucose is absorbed into the blood, eliciting an insulin response. The digestion and absorption kinetics (rapid or slow) depends on starch structure.

**OBJECTIVE:** To study the relationship between the *in vivo* glycemic and insulinemic index and the *in vitro* digestibility characteristics of six bakery products, made from non-conventional wholemeal/wholegrain flours.

**METHODS:** We analyzed *in vitro* the rapidly- and slowly- available glucose (RAG and SAG), the rapidly- and slowly- digestible starch (RDS and SDS), and the resistant starch (RS) fraction of the six wholemeal/wholegrain products and one white type of bread. The glycemic and the insulinemic index (GI and II respectively) were estimated by *in vivo* testing in a group of eleven healthy individuals.

**RESULTS:** The GI of the wholemeal/wholegrain flour biscuits and breads were low, (range 28±3.2 to 41±3.9, Mean±SEM) correlating with the II. RAG positively correlated with both GI and II, with fiber having a marginal correlation.

**CONCLUSIONS:** Our findings indicate that both conventional and non-conventional wholemeal/wholegrain bakery products have low GI and moderate II, correlating to *in vitro* starch digestibility and the type of processing.

## 1. Introduction

Carbohydrates are a common constituent of the human diet, representing 50% of daily energy intake in the Western diet [1]. In a typical diet, carbohydrates are mostly available as starch. Starch is digested in the human small intestine in order to deliver glucose that is absorbed into the blood. The rate and the extend of the digestion of the carbohydrate consumed is influenced by its botanical origin as this determines the amylose:amylopectin ratio and the structure of the starch granule [2]. Furthermore, the macronutrient content (fat, fiber, protein etc.) of the diet strongly influences the rate of digestion and the subsequent physiological responses to glucose absorption manifested as glycemia and insulinemia [3]. Food processing is also important, as this determines the extend of starch gelatinization, particle size and the integrity of plant cell wall [4]. Therefore, different cereal-based bakery products exhibit a wide variation in GI and II, optional to phytochemical characteristics, the accompanying nutrient composition as well as the particular processing.

Dietary carbohydrates can be classified according to their chemical composition and likely site, rate and extend of digestion [5]. Starch is therefore classified as rapidly digestible starch (RDS), slowly digestible starch (SDS), and resistant starch (RS). Similarly, rapidly available glucose (RAG) and slowly available glucose (SAG), reflect the rate at which glucose (from sugars and starch, including maltodextrins) becomes available for absorption in the human small intestine and appears in the blood. Both RAG and SAG values strongly influence the glycemic index of cereal-based foods [6].

The glycemic index (GI) has been introduced in order to classify carbohydrate foods based on how they influence postprandial plasma glucose response. GI is expressed as the percentage increase in the glucose area under the curve of a test food against a standard food such as glucose or white bread [7]. Apart from the food composition, the GI is also influenced by the interplay of post-absorptive metabolic events wherein insulin has the most important contribution. Therefore, the GI is closely related to the insulinemic index (II) as insulin is secreted in the blood in direct response to the postprandial increase in blood glucose and coordinates both glucose metabolic handling as well as clearance of any excess from the circulation. The II can be determined in tandem with the corresponding GI for a given test food with concomitant measuring of insulin as well as glucose levels in the same blood sample. Many types of food have been classified in terms of their glycemic load. It is inevitable that diets containing large amounts of rapidly digested carbohydrates, apart from causing high glycemia, they also put a challenging pressure upon the physiological processes of glucose homeostasis by eliciting a dramatic and prolonged insulin response. Chronic exposure to such a condition (in combination with the host lifestyle and sedentary habits), encourages obesity and increases the relative risk for type 2 diabetes mellitus, particularly correlating the combination of high glycemic load with low cereal fiber intake [8, 9]. On the other hand, diets rich in slowly digested carbohydrates and high in fiber, offer considerable protection against metabolic disease [10]. Therefore, there is a universal demand of cereal products that combine high energy and nutrient merit with low glycemic load [11, 12]. Bakery goods (biscuits and bread) are popular complements of a typical Mediterranean diet. Particularly those made of non-conventional, non-processed flour mixes, enriched with whole grain cereals and plant protein, may combine the benefits of high energy, high nutritive value and low glycemic load.

The aim of the present study was to test the glycemic and insulinemic effect of particular biscuits made from non-processed wholemeal and wholegrain flour mixes (consisting of whole grain spelt, whole grain oat, barley and lupine flour), as well as three types of wholemeal/wholegrain bread. White bread was used as reference. The biscuits and bread were tested using a modification of the method of Englyst et al [13] so as to determine their SDS, RDS, RS, RAG and SAG. By using this *in vitro* digestion assay and estimating the corresponding glycemic and insulinemic index in a group of healthy volunteers, we studied the effect of composition (wholemeal/wholegrain vs processed flour) and glucose mobilization kinetics (RAG vs SAG) on the postprandial glycemic and insulinemic response.

## 2. Materials, subjects and methods

### 2.1 Test bakery products

Biscuit 1 (“Breakfast Belvita”cinnamon/brown sugar – NABISCO Mondelez Global LLC, East Hanover HJ 07936, USA. Biscuit 2 “Olympic Runner” cinnamon/tahini and Biscuit 3 “Athenian Philosopher” fig/anise were both made by OLYRA S.A., Alexandroupolis, Evros 68133 Greece P.O.B.:1095, https://olyrafoods.com/). All three wholemeal and the white-type bread flour mixtures and the breads were prepared and baked at the premises of OLYRA FOODS from flour mixes made by FLOURMILLS THRAKIS – I. OUZOUNOPOULOS S.A., Alexandroupolis, Evros, Greece.

Pepsin (from porcine gastric mucosa) 0.7 FIP-U/mg and pancreatin (from porcine pancreas) 350FIP-U/g protease, 6,000 FIP-U/g lipase, 7,500FIP-U/g amylase were from Merck KGaA, Darmstadt, Germany. Amyloglucosidase (from *Rhizopus sp*.) 5,000U; 42U/mg and Invertase, 300U/mg were from Megazyme, BRAY BUSINESS PARK, Southern Cross Rd, Bray, Co. Wicklow, A98 YV2, Ireland.

### 2.2 In vitro measurement of free sugar glucose and fructose, RAG, SAG, total glucose and starch fractions of bakery products

Samples of bakery products (containing <0.6 g carbohydrate) were weighted to the nearest milligram into 50-ml polypropylene centrifuge tubes. Free sugar glucose and fructose, RAG, SAG, total glucose and starch fractions (RDS, SDS and RS contents) were analyzed by the protocol of Englyst et al. [13]. Free sugar glucose and fructose, as well as glucose portions taken at 20-,120- and 240-min endpoints (G_20_, G_120_ and G_240_ respectively), were collected in absolute ethanol and quantified by HPLC in a private laboratory (Q & Q Analysis, Food quality services laboratory, Mikrasiaton 67 & Kountouriotou, 38333, Volos, Greece) by standard validated analytical protocols utilized therein.

### 2.3 Subjects

Eleven healthy volunteers, (6 men and 5 women aged 40 ± 20 years with body mass index (BMI) of 22.9 ± 1.9 took part in the study. None of them had any history of metabolic disease, and they were not taking any drugs that might have influenced their metabolic profile. Ethical permission for the study was obtained from the Democritus University of Thrace Ethical Committee. On test days, after an overnight fast following a day on which they were advised to abstain from alcohol and excessive exercise, they arrived at the laboratory and an iv cannula was inserted into the antecubital vein. A fasting blood sample was taken before the test bakery product was provided. It was eaten within 10 minutes and subsequent blood samples were taken at 15, 30, 60, 90 and 120 minutes after consumption, into vacutainers without anticoagulants and were allowed to clot, immediately centrifuged, and kept at −80°C till analysis. All volunteers were tested in sequence consuming one product at a time (the order was randomized and different for each subject), with intervals of ≥ 1 week between each test. Portion sizes were 60g for the four types of bread. For the biscuits, 50g of Biscuit 1 and 37,5g of Biscuits 2 and 3 were administered as individually packed «suggested portion» by the respective manufacturers. The volunteers also received 30g of soluble pure glucose (Merck) as a reference experiment in order to assess the area under the curve AUC under optimally available «free» glucose. Blood insulin was measured by RIA (I-125 IRMA Kit REF: RK-400CT, by IZOTOP, Institute of Isotopes Ltd, H-1535 Budapest, P.O.B.851 Hungary). Glucose was measured in a SIEMENS ADVIA automatic analyzer using a Medicon Kit (REF: 1417-0017, MEDICON HELLAS AE, Greece).

### 2.4 Calculations and statistics

RAG, SAG and the various starch fractions were calculated by the formulae detailed by Englyst et al [13] as follows:

RAG = G_20_

SAG = G_120_ - G_20_

RDS = (G_20_ - FSG) x 0.9 SDS = (G_120_ – G_20_) x 0.9

Total starch = (total glucose - FSG) x 0.9

Resistant starch = (total glucose – G_120_) x 0.9

The glycemic and the insulinemic responses were calculated as the incremental areas under the respective curves above the fasting levels, ignoring any area below the fasting value.

All statistical analyses were performed using the R programming language version 4.1.2 [14]. Linear regression was performed using the base R lm command. p values<0.05 were considered statistically significant. Graphics were drawn using the ggplot2 package (version 3.3.5).

## 3. Results

### 3.1 Glycemic and insulinemic indices

The nutrient profile (g/portion used in the glycemic index test) of the seven selected bakery products is shown in Table 1. The content of the *in vitro* digestion fractions (g/100 g of product), GI and II data for each product are shown in Table 2. Mean RAG content was 9.48 ± 1.41 g per 100 g product for the three wholemeal/wholegrain biscuits and bread; the white bread had a RAG value of 45.5 g per 100g of product. Mean SAG was 8.48 + 2.41g per 100 g product for the three wholemeal/wholegrain biscuits and bread; the white bread had a SAG value of 0.8 g per 100g of product.

**TABLE 1.** Nutritional composition of the test bakery products.

|  | BISCUIT 1<br>"BREAKFAST<br>BELVITA®" | BISCUIT 2<br>"OLYMPIC<br>RUNNER®" | BISCUIT 3<br>"PHILOSOPHER®" | WHOLEMEAL<br>BREAD 1 | WHOLEMEAL<br>BREAD 2 | WHITE BREAD<br>3 | WHOLEMEAL<br>BREAD 4 |
| --- | --- | --- | --- | --- | --- | --- | --- |
| Product Supplier | NABISCO | OLYRA FOODS | OLYRA FOODS | OLYRA FOODS | OLYRA FOODS | OLYRA FOODS | OLYRA FOODS |
| Portion size (g) | 50 | 37.5 | 37.5 | 60 | 60 | 60 | 60 |
| Energy in 100g (kCal) | 436 | 461 | 469 | 416.86 | 388.5 | 347 | 416.8 |
| Protein in 100g | 7.8 | 10.4 | 9.1 | 16.1 | 14.75 | 10.5 | 16.3 |
| Carbohydrate in 100g | 66 | 57 | 61.4 | 54.8 | 54.65 | 73.3 | 55.7 |
| Sugars in 100g | 20 | 22.43 | 20.1 | 2.35 | 2.5 | 0.5 | 2.24 |
| Fiber in 100g | 6.8 | 9.7 | 5.6 | 8.45 | 5.5 | 1.5 | 8.9 |
| Fat in 100g | 14 | 19.1 | 19.1 | 5.25 | 11.2 | 1.4 | 1.33 |
| Saturated Fat in 100g | 1.5 | 7.9 | 8.5 | 0.686 | 1.6 | 0.4 | 0.686 |
| Energy / portion (kCal) | 218 | 172.9 | 175.9 | 250.1 | 233.1 | 208.2 | 250.1 |
| Protein / portion | 3.9 | 3.9 | 3.4 | 9.7 | 8.85 | 6.3 | 9.8 |
| Carbohydrate / portion | 33 | 21.4 | 23.02 | 32.9 | 32.8 | 43.98 | 33.42 |
| Sugars / portion | 10 | 8.4 | 7.53 | 1.41 | 1.5 | 0.3 | 1.34 |
| Fiber / portion | 3.4 | 3.64 | 2.1 | 5.1 | 3.3 | 0.9 | 5.34 |
| Fat / portion | 7 | 7.2 | 7.16 | 3.15 | 6.72 | 0.84 | 0.8 |
| Saturated Fat / portion | 0.75 | 2.95 | 3.2 | 0.41 | 0.96 | 0.24 | 0.41 |
| Rolled Oats, Rye Flakes/100g | 38 | - | - | - | - | - | - |
| Wholegrain flour (Spelt, oat, barley, lupin)/100g |  | 20 | 20 | - | - | - | - |
| Sunflower kernels, linseed, pea fiber, sesame seeds, salt, wheat bran, malt flour (barley, wheat), malt extract (barley). Dried sourdough* (in 100g) | - | - | - | 21.1 | 21.1 | - | - |
| Sunflower kernels, linseed, sesame seeds, oat flakes, rye bran, wheat bran, Dried sourdough* (in 100g) | - | - | - | - | - | - | 39.0 |

(\* contains components from wheat, rye and barley.)

**TABLE 2.**
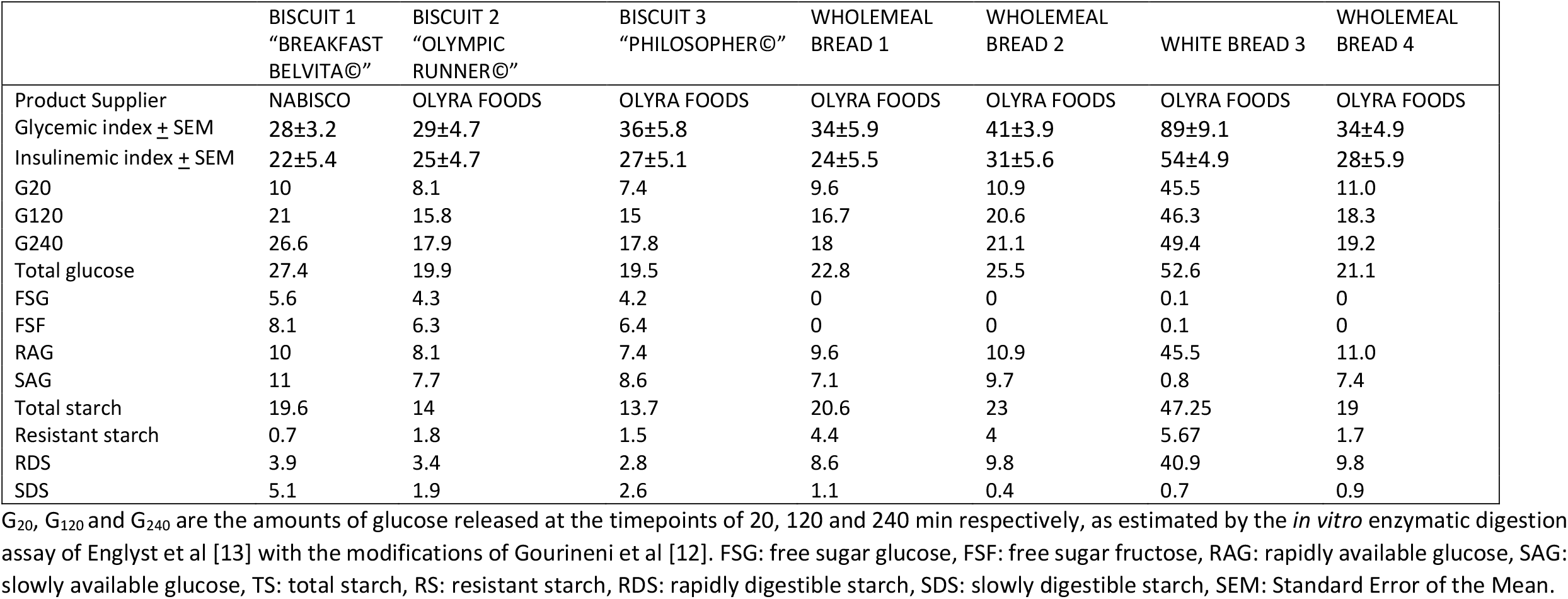
Glycemic and insulinemic indices and starch digestibility parameters (g/100 g of product) of the test bakery products.

|  | BISCUIT 1<br>"BREAKFAST<br>BELVITA®" | BISCUIT 2<br>"OLYMPIC<br>RUNNER®" | BISCUIT 3<br>"PHILOSOPHER®" | WHOLEMEAL<br>BREAD 1 | WHOLEMEAL<br>BREAD 2 | WHITE BREAD 3 | WHOLEMEAL<br>BREAD 4 |
| --- | --- | --- | --- | --- | --- | --- | --- |
| Product Supplier | NABISCO | OLYRA FOODS | OLYRA FOODS | OLYRA FOODS | OLYRA FOODS | OLYRA FOODS | OLYRA FOODS |
| Glycemic index ± SEM | 28±3.2 | 29±4.7 | 36±5.8 | 34±5.9 | 41±3.9 | 89±9.1 | 34±4.9 |
| Insulinemic index ± SEM | 22±5.4 | 25±4.7 | 27±5.1 | 24±5.5 | 31±5.6 | 54±4.9 | 28±5.9 |
| G20 | 10 | 8.1 | 7.4 | 9.6 | 10.9 | 45.5 | 11.0 |
| G120 | 21 | 15.8 | 15 | 16.7 | 20.6 | 46.3 | 18.3 |
| G240 | 26.6 | 17.9 | 17.8 | 18 | 21.1 | 49.4 | 19.2 |
| Total glucose | 27.4 | 19.9 | 19.5 | 22.8 | 25.5 | 52.6 | 21.1 |
| FSG | 5.6 | 4.3 | 4.2 | 0 | 0 | 0.1 | 0 |
| FSF | 8.1 | 6.3 | 6.4 | 0 | 0 | 0.1 | 0 |
| RAG | 10 | 8.1 | 7.4 | 9.6 | 10.9 | 45.5 | 11.0 |
| SAG | 11 | 7.7 | 8.6 | 7.1 | 9.7 | 0.8 | 7.4 |
| Total starch | 19.6 | 14 | 13.7 | 20.6 | 23 | 47.25 | 19 |
| Resistant starch | 0.7 | 1.8 | 1.5 | 4.4 | 4 | 5.67 | 1.7 |
| RDS | 3.9 | 3.4 | 2.8 | 8.6 | 9.8 | 40.9 | 9.8 |
| SDS | 5.1 | 1.9 | 2.6 | 1.1 | 0.4 | 0.7 | 0.9 |

G<sub>20</sub>, G<sub>120</sub> and G<sub>240</sub> are the amounts of glucose released at the timepoints of 20, 120 and 240 min respectively, as estimated by the *in vitro* enzymatic digestion assay of Englyst et al [13] with the modifications of Gourineni et al [12]. FSG: free sugar glucose, FSF: free sugar fructose, RAG: rapidly available glucose, SAG: slowly available glucose, TS: total starch, RS: resistant starch, RDS: rapidly digestible starch, SDS: slowly digestible starch, SEM: Standard Error of the Mean.

In Figures 1 and 2, the glycemic and insulinemic response for the seven test foods compared to the 30 grams of glucose administered are shown, respectively. In Table 2, the GI and II of the test foods are shown. The mean GI of the wholemeal flour biscuits and breads was 33.7 with a range from 28 to 41, with the white bread having a GI of 89±9.1. The mean II of the wholemeal biscuits and bread was 21 with a range from 22 to 31, with the white bread having an II of 54±4.9.

**Figure 1.**
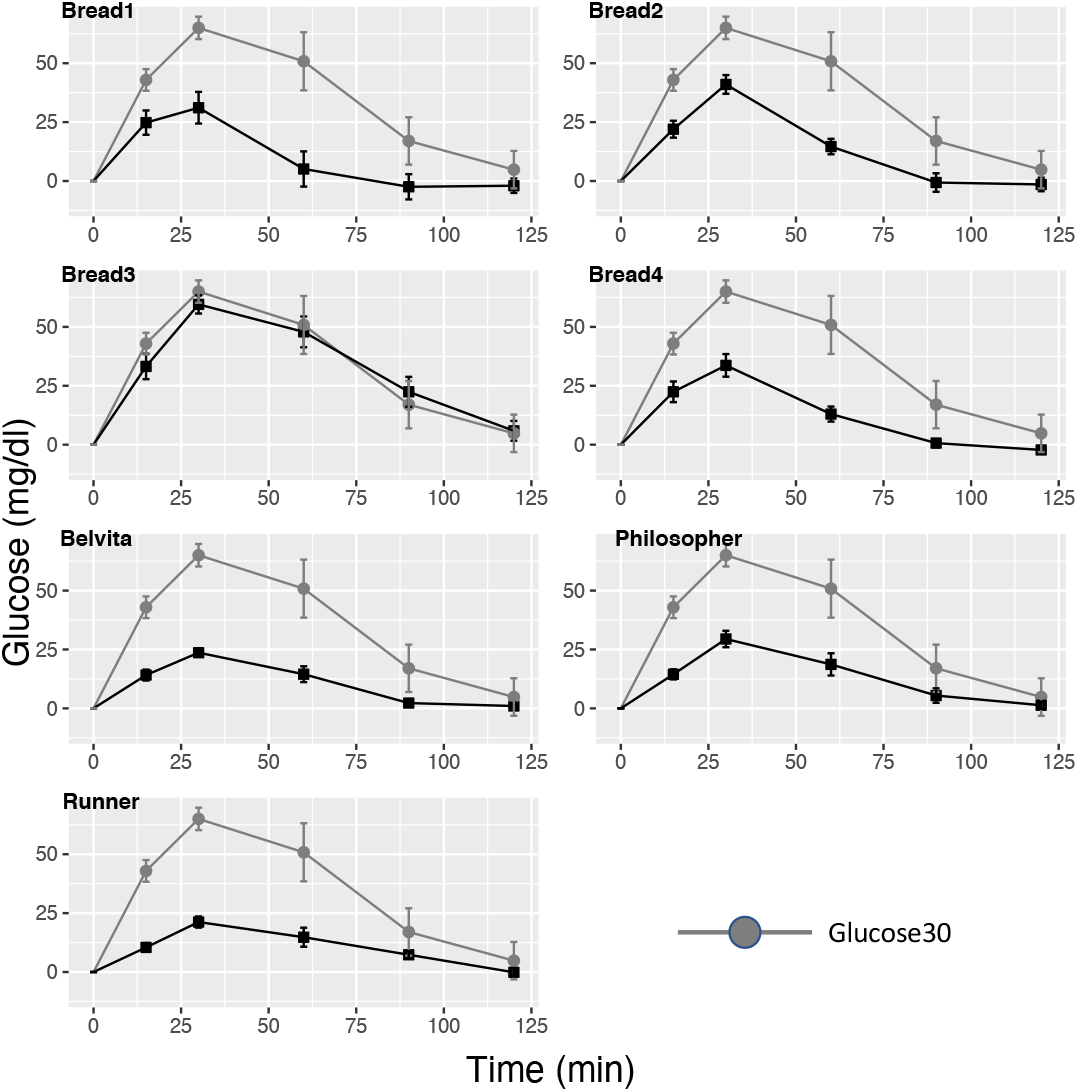
Glycemic response (± SEM increase in blood glucose) to the 7 test foods. The glucose response after oral administration of 30 gr glucose is shown in all subgraphs. For comparison purposes, all x and y axes ranges are the same.

**Figure 2.**
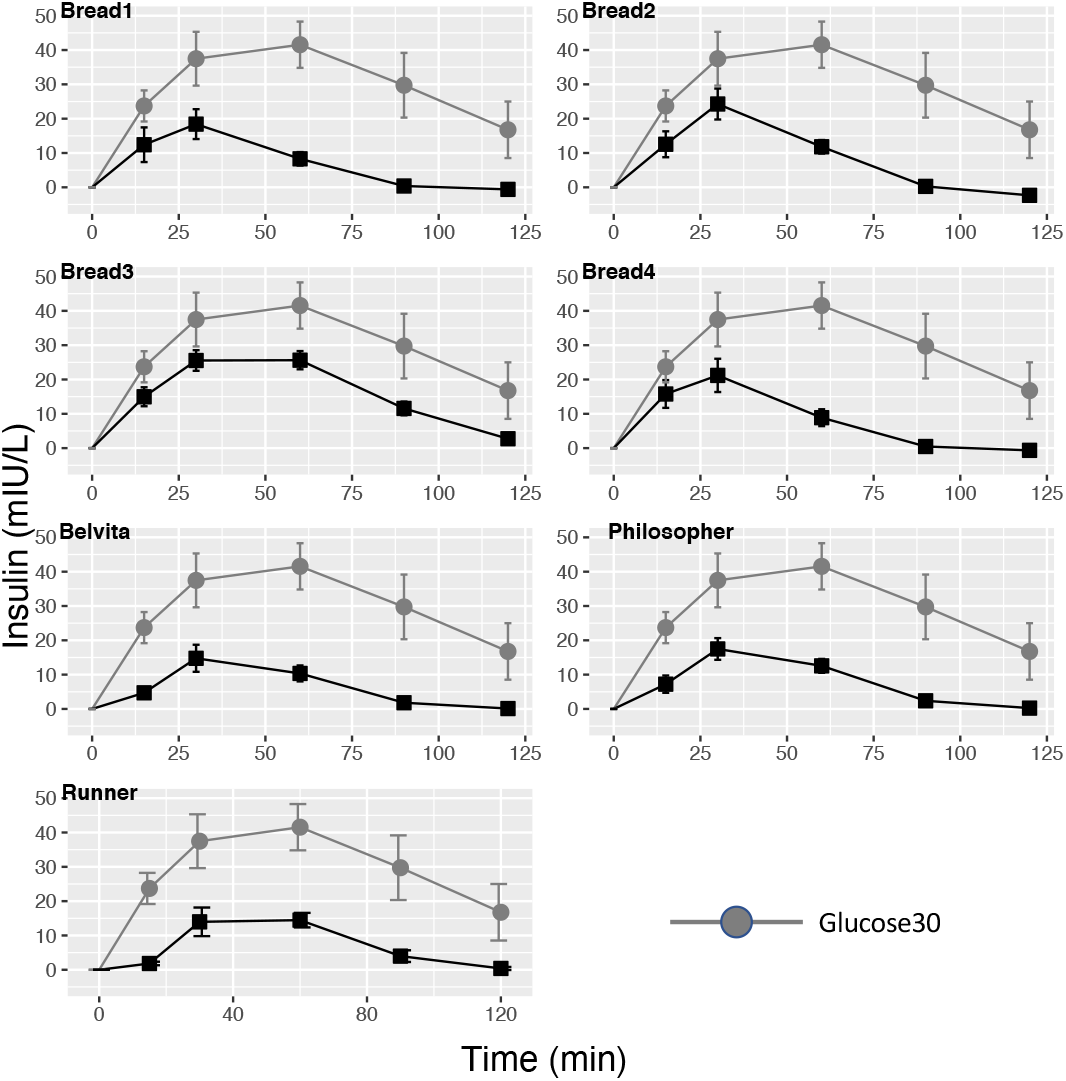
Insulinemic response (± SEM increase in serum insulin) to the 7 test foods. The glucose response after oral administration of 30 gr glucose is shown in all subgraphs. For comparison purposes, all x and y axes ranges are the same.

### 3.2 Linear regression for the glycemic index dependence on RAG, protein and fiber content

Various linear regression models were constructed to determine the dependence of the GI upon food RAG, protein and fiber content. For these models, one extreme outlier (with a glycemic response near 0) was excluded. The results are shown in Table 3.

**TABLE 3.** Dependence of the Glycemic Index on RAG, fiber and protein content.

| Model number | Model | R<sup>2</sup> | F statistic | F statistic<br>p | Intrcpt±SE | P(Intrcpt≠0) | RAG<br>coef±SE | P(RAG<br>coef≠0) | Fiber<br>coef±SE | P(Fiber<br>coef≠0) | Protein<br>coef±SEM | P(Protein<br>coef≠0) |
| --- | --- | --- | --- | --- | --- | --- | --- | --- | --- | --- | --- | --- |
| 1 | $GI = \beta_0 + \beta_1 \cdot RAG$ | 0.952 | F(1,5)=120.8 | 0.0001 | 19.2±2.7 | 0.0008 | 1.5±0.14 | 0.0001 | | | | |
| 2 | $GI = \beta_0 + \beta_1 \cdot fiber$ | 0.288 | F(1,5)=3.4 | 0.1234 | 71.2±17.4 | 0.0094 | | | -8.7±4.72 | 0.1234 | | |
| 3 | $GI = \beta_0 + \beta_1 \cdot protein$ | -0.191 | F(1,5)=0.04 | 0.8526 | 37.3±23.4 | 0.172 | | | | | 0.6±3.32 | 0.8526 |
| 4 | $GI = \beta_0 + \beta_1 \cdot RAG + \beta_2 \cdot fiber$ | 0.941 | F(2,4)=48.6 | 0.0016 | 18.0±8.7 | 0.1075 | 1.6±0.21 | 0.0017 | 0.3±1.82 | 0.8922 | | |
| 5 | $GI = \beta_0 + \beta_1 \cdot RAG + \beta_2 \cdot protein$ | 0.945 | F(2,4)=52.1 | 0.0014 | 16.6±5.4 | 0.0378 | 1.5±0.15 | 0.0005 | | | 0.4±0.72 | 0.6113 |
| 6 | $GI = \beta_0 + \beta_1 \cdot fiber + \beta_2 \cdot protein$ | 0.564 | F(2,4)=4.9 | 0.0847 | 56.4±15.5 | 0.0218 | | | -14±4.51 | 0.036 | 5±2.45 | 0.1111 |
| 7 | $GI = \beta_0 + \beta_1 \cdot RAG + \beta_2 \cdot fiber + \beta_3 \cdot protein$ | 0.930 | F(3,3)=27.7 | 0.0109 | 19.9±9.9 | 0.1382 | 1.4±0.3 | 0.0183 | -1.4±3.24 | 0.7025 | 0.8±1.32 | 0.5722 |

GI: Glycemic Index, Intrcpt: Intercept, coef: Coefficient, SEM: Standard Error of the Mean, RAG: Rapidly Available Glucose

The best model (as determined by the R^2^ value) was that for the dependence of the GI only upon RAG (model number 1). The R^2^ was 0.952, with F(1,5)=120.8, p= 0.0001. The intercept β_0_ was 19.2, with a standard error of 2.7 (p=0.0008) and the slope β_1_ was 1.5 with a standard error of 0.14 (p=0.0001).

The R^2^ for the dependence of the GI on either fiber or protein content (models 2 and 3) were low (0.288 and −0.191, respectively), with the respective fiber and protein coefficients not being statistically significantly different from 0.

When the linear dependence of the GI upon RAG and either fiber or protein was tested (models 4 and 5), the R^2^ values were good (0.941 and 0.945, respectively). However, neither the coefficient for fiber content (model 4), nor the one for protein content (model 5), were statistically significantly different from 0, whereas the RAG coefficient remained statistically significantly different from 0 and with almost the same values as in model 1, having RAG as the only independent variable (1.5±0.14 for RAG only, 1.6±0.21 for RAG plus fiber and 1.5±0.15 for RAG plus protein).

When the RAG was not included as an independent variable and the GI dependence upon only fiber and protein was tested (model 6), the R^2^ value was moderately good (0.564), but much lower than the R^2^ for RAG only. Furthermore, the coefficient for fiber was of marginal statistical significance (p=0.036), whereas the protein dependence was not statistically significant (p=0.1111).

Finally, when all independent variables (RAG, fiber and protein) were included in the model (model 7), the R^2^ value was very good (0.930). However, only the RAG dependence remained statistically significant (p=0.0183) and in the same range as in models 1, 4 and 5 (1.4±0.30), whereas the fiber and protein coefficients were not statistically significant.

When, instead of RAG, SAG was used as an independent variable, in none of the models was there a statistically significant SAG coefficient. When both RAG and SAG were used, only the coefficient for RAG was statistically significant (data not shown).

### 3.3 Linear regression for the insulinemic index dependence of RAG, protein and fiber content

For the linear regression models described below, two extreme outliers with insulinemic response ≅ 0 were excluded.

Dependence of the insulinemic index on RAG, fiber or protein content, dovetails with that of the glycemic index, as shown in Table 4. The best model was again the one with RAG (model 1) as the only independent variable (R^2^=0.922), with intercept=18.9±1.8 and a coefficient for RAG=0.8±0.09, with both being statistically significant different from 0.

**TABLE 4.**
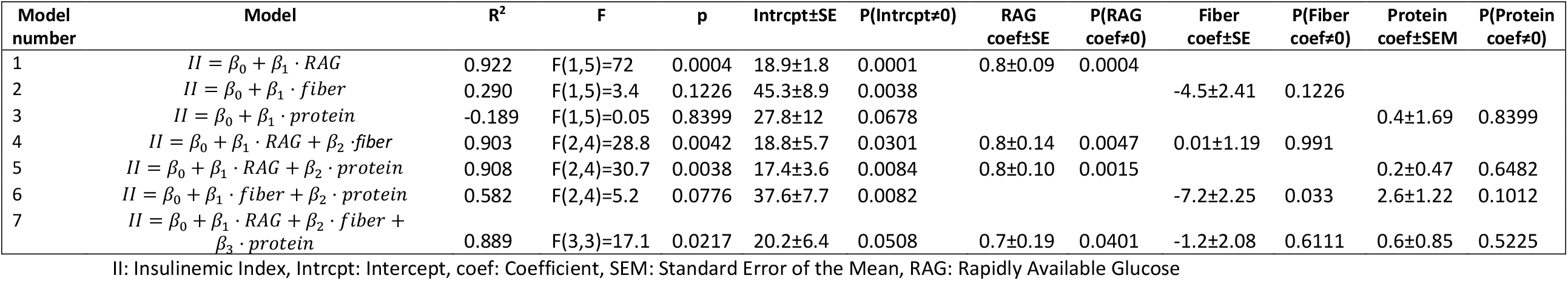
Dependence of the Insulinemic Index on RAG, fiber and protein content.

When either fiber or protein were the sole independent variables (models 2 and 3), the R^2^ was not good when compared to model 1 (0.290 and −0.189, respectively). Also, both the intercepts and the linear coefficients were not statistically significant different from 0.

When the RAG and either fiber or protein were used as independent variables (models 4 and 5), the R^2^ was good (0.903 and 0.908, respectively), but neither the fiber coefficient in model 4, not the protein coefficient in model 5, were statistically significantly different from 0. On the other hand, the linear coefficient for RAG in all three models was essentially the same (0.8±0.09 for model 1, 0.8±0.14 for model 4 and 0.8±0.10 for model 5). In all three models, the RAG coefficients were statistically significant, although for model 4 and 5 the significance was reduced.

When fiber and protein were used together as independent variables (model 6), but in the absence or RAG, the R^2^ (although satisfactory, with a value of 0.582), was much less than that of model 1, and only the fiber coefficient was of marginal statistical significance.

Finally, when all RAG, fiber and protein were included in the model as independent variables (model 7), the R^2^ was good (0.889), but neither the fiber nor the protein coefficients were statistically significant. Only the linear coefficient for RAG remained marginally significant, and in the same range as in models 1, 4 and 5 (0.7±0.19).

As with the glycemic index, when SAG was used as the independent variable instead of RAG, again no statistically significant SAG coefficient was evident. When both RAG and SAG were included, only the RAG coefficient was statistically significant (data not shown).

### 3.4 Glycemic index dependence of the Insulinemic index

A linear regression model was created with the glycemic index as the independent parameter and the insulinemic index as the dependent one. The graph of the linear regression line is shown in Figure 3.

**Figure 3.**
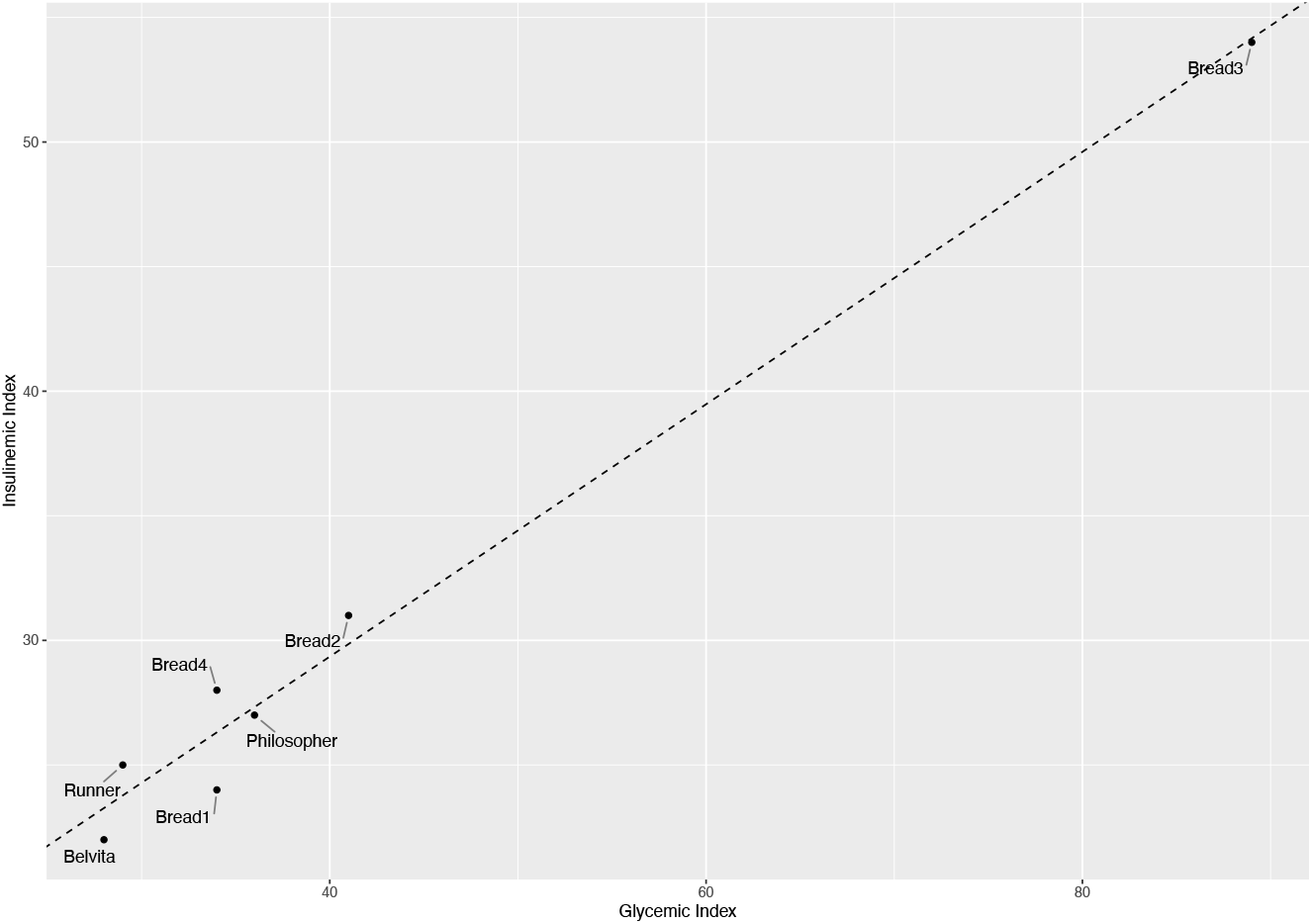
Scatterplot and linear regression line for the dependence of the insulinemic upon the glycemic index.

A strong linear dependence of the insulinemic upon the glycemic index is evident. The R^2^ was 0.979, F(1,5)=275.5, p<0.00001. The intercept was 9.09±1.40 (p=0.0015) and the slope was 0.51±0.03 (p<0.00001).

## 4. Discussion

In the present study, we analyzed six bakery products (three biscuits and three types of bread) made from non-conventional flour mixtures. The biscuits contained whole grain spelt, whole grain oat, barley and lupine flour and the breads contained pregelatinized wheat flour, sunflower kernels, linseed, pea fiber, sesame seeds, wheat bran, malt flour from barley and wheat, malt extract from barley as well as dried sourdough (Table 1). We also analyzed one white type of bread, as a reference example containing a conventionally processed flour mixture. We applied *in vitro* and *in vivo* testing methodology as well as a number of statistical models in order to study the influence of inherent food properties (glucose content, type of starch, the presence of other macronutrients (protein, fiber) and different food processing) upon both the Glycemic and Insulinemic index.

To this end, we analyzed the SDS, RDS and RS fractions of the test bakery products by an *in vitro* assay measuring the amount of glucose released at different time points during incubation with digestive enzymes under standardized conditions [13]. We also determined both the GI and II as *in vivo* reporters of the direct glycemic response to these foods upon consumption.

In line with previous reports, we found that the RAG content strongly affects both the GI and the II [15]. Although there were no significant differences in the RAG fraction among the three biscuits and the three types of wholemeal/wholegrain bread tested (9.48 ± 1.41 g of glucose /100g of product), the RAG estimated for the white bread by a value of 45.5 was by far the highest. Similarly, both the GI and II for the bread made from white processed flour were much higher (89±9.1 and 54±4.9 respectively) compared to those for the wholemeal/wholegrain products.

From the application of a variety of linear regression models that were tested for the glycemic (Table 3) and insulinemic (Table 4) indices, it can be concluded that the main factor affecting both the glycemic and the insulinemic index is the rapidly available glucose (RAG).

In the case of the glycemic index, the R^2^ was 0.952, which means that variation in RAG explains 95.2% of the GI variation. The linear coefficient for RAG was 1.5±0.14, meaning that (under the conditions of this study), an increase of 1 unit in RAG, correlates with an increase of the glycemic index by 1.5±0.14 units. The RAG was the only independent variable found to be correlated to the glycemic index in a statistically significant manner.

For both GI and II, when SAG was used in the place of RAG, no statistically significant correlation was observed. This was also the case when both RAG and SAG were included. This shows that the main determinant of both the GI and the II is the glucose that is rapidly available for absorption from the digestive tract (glucose available in the first 20 minutes in the *in vitro* digestion) and not the more slowly available glucose (glucose released between 20 and 120 min in the *in vitro* digestion).

The only other coefficient of marginal statistical significance was that for fiber, when used in a model with fiber and protein together as independent variables. In this case the coefficient for fiber was −14±4.51, meaning that an increase of 1 unit in fiber content, correlates with a decrease of the glycemic index by 14 units. However, this model explained only 58.2% of the GI variation, as opposed to 95.2%, with RAG being the most prominent independent variable.

Similar correlations as in the case of the glycemic index were also observed in the case of the insulinemic index. Variation in RAG alone explained 92.2% of variation in the insulinemic index, with a 1 unit increase in RAG being correlated with an increase of 0.8±0.09 units in the insulinemic index. When fiber and protein were included as independent variables in the absence of RAG, the model explained 58.2% of the insulinemic index variation, with a decrease of 7.2±2.25 units for a 1 unit increase in fiber, but again being of marginal statistical significance.

Interestingly, both the observed glycemic and insulinemic indices were marginally dependent on fiber content when it was considered as an independent variable. Therefore, our findings support the current knowledge that mainly RAG and the presence of fiber certainly highlight a major difference in nutritional composition between wholemeal/wholegrain and processed white flour in the bakery products tested. Wholemeal flour still contains all the natural fiber available in the entire kernel of the grain, by contrast to white flour that contains mostly the endosperm portion of the grain kernel with germ and bran having been removed. Such being the case for the white bread flour, baking in high humidity aids gelatinization and alters dramatically the properties of starch, making it more readily available to enzymic digestion [4]. It is therefore fair to say that fiber may not *per se* strongly affect GI and II as an independent variable, nevertheless the consequences of its removal may indirectly reflect on both indices by augmenting the rate of starch digestion (that is essentially RAG).

Furthermore, the combination of wholemeal/wholegrain flour mixtures and sourdough in the three wholemeal/wholegrain bread tested, very possibly explain their low GI and II observed [16].

For all products tested, when the linear dependence of the II upon the GI was examined (shown in Figure 3), the resulting R^2^ was 0.979, meaning that the variation in GI explains 97.9% of the II variation. Also, the slope was 0.51±0.03, meaning that for a 1 unit increase in the GI, the II increases by approximately 0.5 units.

## 5. Conclusion

To the best of our knowledge, there are limited studies that correlate the *in vitro* digestibility profile of foods with both the glycemic and the insulinemic index. Taken together, GI and II can provide an important insight into the post-absorptive glycemic physiology. Our observations provide further evidence that the RAG content of bakery products is the most prominent independent variable that correlates with both the glycemic and insulinemic response and characterizes test foods with respect to their glycemic properties.

Finally, bakery products made from both conventional and non-conventional wholemeal/wholegrain flour mixtures have a low GI and a moderate II. By being a staple part of the diet, they may present a healthier alternative to those made with processed flour by offering protection against metabolic disease.

## List of abbreviations

GI: Glycemic Index
II: Insulinemic Index
RAG: Rapidly Available Glucose
SAG: Slowly Available Glucose
RDS: Rapidly Digestible Starch)
SDS: Slowly Digestible Starch (SDS)
RS: Resistant Starch
FSG: Free Sugar Glucose
SEM: Standard Error of the Mean

## Author Contributions

- Charalampos Papadopoulos: Conception; Performance of work, (*in vitro/in vivo* study); writing the article
- Constantine Anagnostopoulos: Conception; performance of work; interpretation of data; writing the article
- Athanasios Zisimopoulos: Conception; performance of work
- Maria Panopoulou: Conception; performance of work
- Dimitrios Papazoglou: Conception; performance of work; interpretation of data
- Anastasia Grapsa: Performance of work, (*in vivo* study)
- Thaleia Tente: Performance of work, (*in vitro/in vivo* study)
- Ioannis Tentes: Conception; performance of work; interpretation of data; writing the article

